# The transcriptional and translational outcomes for pseudogenes in bacterial endosymbionts

**DOI:** 10.64898/2026.02.04.703838

**Authors:** Arkadiy Garber, Justus Nwachukwu, Ryan Stikeleather, Courtney York, John P. McCutcheon

## Abstract

Intracellular bacteria in the early stages of host adaptation often show extraordinarily disrupted genomes, where up to half of their ancestral genes are found in a pseudogenized state. The mealybug *Pseudococcus longispinus* hosts two bacterial endosymbionts with high pseudogene loads, *Symbiopectobacterium endolongispinus* and *Sodalis endolongispinus*. Here, we measure transcript abundance, ribosome-associated RNA, and protein abundance in these bacterial symbionts to understand how bacteria avoid (or fail to avoid) accumulating large amounts of non-functional RNAs and proteins from these pseudogenes. Consistent with previous work, we show that pseudogene transcripts remain detectable, but at lower levels compared to those from intact and functional genes, and that relatively few pseudogenes yield detectable proteins in proteomic data. However, we find that many pseudogene transcripts still bind to *Symbiopectobacterium* ribosomes, and uncover a possible role for the tmRNA ribosome rescue system in the targeting of pseudogene proteins for degradation. Our results suggest a possible mechanism by which bacterial endosymbionts remove aberrant pseudogene-derived proteins during the critical time when many pseudogenes have formed but not enough time has passed for sequence evolution to erode ribosome binding sites from pseudogene transcripts.

**Significance:** Bacteria transitioning from free-living to host-dependent lifestyles often go through a transitory period where large numbers of genes are broken but not yet deleted. How cells navigate this period without producing useless or toxic gene products remains poorly understood. By combining transcriptomic, proteomic, and ribosomal profiling data from two bacterial endosymbionts, we uncover a possible mechanism that cells use to mitigate the presence of thousands of newly formed pseudogenes. We show that pseudogenes are still widely represented in the transcriptome and are often associated with ribosomes, but rarely yield measurable protein products. RNA sequencing from purified ribosomes suggests that the tmRNA ribosome rescue system may act as a short-term quality control mechanism during early stages of genome reduction. These findings provide a mechanistic glimpse into how endosymbionts survive the unstable phase between gene inactivation and gene deletion, a fleeting but critical window in the evolution of endosymbiosis.

## INTRODUCTION

Contrasting their metabolic, phylogenetic, and environmental diversity (Eren and Banfield, 2024), bacteria have remarkably stable and predictable genome structures. Bacterial genomes tend to retain few broken or inactivated genes (also known as pseudogenes), and encode about one functional gene every 1,000 base pairs (bp) (Kirchberger et al., 2020). Reductions from this uniform gene density—that is, fewer functional genes per region of the genome—are rare, and are mostly found in organisms that have recently undergone shifts in their environment, such as in bacteria that have recently transitioned from a free-living to an intracellular state. Living inside a eukaryotic cell brings with it strong environmental and population genetic forces that make large numbers of genes redundant (Boyd et al., 2024). While the tiniest bacterial genomes, such as those found in endosymbionts of sap-feeding insects (McCutcheon and Moran, 2011), show the typical high functional gene density, the path taken to arrive at this compact state includes transitional periods where the proportion of pseudogenes, relative to functional genes, is very high (Toh et al., 2006; Burke and Moran, 2011; Oakeson et al., 2014; Nechitaylo et al., 2021). It is now known that in the early stages of endosymbiosis, when a free-living bacterium transitions to becoming a vertically transmitted symbiont, large numbers of pseudogenes can accumulate. This happens due to functional redundancy with the host cell and other symbionts (Moran et al., 2009; Koga and Moran, 2014), selection against genes that promote a host immune response (Amiel et al., 2010), and a genome-wide reduction in the efficacy of purifying selection (Lerat and Ochman, 2005). In genomes from bacteria that have been caught in a recent transition to the endosymbiotic state, the number of pseudogenes can rival the number of intact genes (Toh et al., 2006; Oakeson et al., 2014). Large numbers of pseudogenes have also been observed in several intracellular bacterial pathogens (Benjak et al., 2017; Cole et al., 2001), likely for similar reasons to those described above for host-beneficial endosymbionts.

How do bacterial cells deal with having hundreds, or even thousands, of broken genes on their genomes? It is reasonable to guess that producing non-functional mRNA would be less problematic than producing non-functional protein products, for both energetic (Lynch and Marinov, 2015) and mechanistic (Kuo and Ochman, 2010) reasons. Previous studies on the transcriptomic and proteomic response to bacterial pseudogene expression support this hypothesis. The tsetse fly endosymbiont *Sodalis glossinidius* shows widespread transcription of its pseudogenes, which number in the thousands (Goodhead et al., 2020). Similar patterns of extensive production of mRNA from pseudogenes were observed in the leprosy bacillus *Mycobacterium leprae* (Suzuki et al., 2006; Williams et al., 2009) and the insect symbiont *Candidatus* Streptomyces philanthi (Nechitaylo et al., 2021). However, most evidence suggests that transcripts produced from pseudogenes do not often lead to the production of measurable protein products. In *M. leprae*, many transcribed pseudogenes lack ribosomal binding sites or start codons, which should reduce the efficiency with which they initiate translation (Williams et al., 2009).

In the insect symbionts *Sodalis glossinidius* and *Candidatus* Streptomyces philanthi, proteomics revealed that relatively few of the numerous pseudogenes found in their transcriptomes were detected as proteins by mass spectrometry (Goodhead et al., 2020; Nechitaylo et al., 2021). However, a survey in *Salmonella enterica* found that about two-thirds of its roughly 160 pseudogenes produced detectable peptides, indicating successful (if often partial) translation of ostensibly inactivated genes (Feng et al., 2022). In *Mycobacterium tuberculosis*, which carries fewer pseudogenes than its leprosy-causing relative, ribosome profiling revealed that numerous pseudogene-derived transcripts are being actively translated (Smith et al., 2022), suggesting that, in some cases, bacteria may not be able to completely stop the translation of non-functional transcripts.

Here, we use the endosymbionts of the long-tailed mealybug, *Pseudococcus longispinus*, to better understand transcriptional and translational outcomes when large numbers of pseudogenes are present on a bacterial genome. *P. longispinus* has an unusual, nested symbiont structure, where two species of recently acquired endosymbionts, *Candidatus* Sodalis endolongispinus (hereafter, *Sodalis endo*.) and *Candidatus* Symbiopectobacterium endolongispinus (hereafter, *Symbiopectobacterium endo*.), reside within the cytoplasm of a long-established endosymbiont called *Candidatus* Tremblaya princeps (hereafter, *Tremblaya princeps*; Husnik and McCutcheon 2016; Garber et al., 2021). Both *Sodalis endo*. and *Symbiopectobacterium endo*. have large genomes containing thousands of pseudogenes, and so are useful models to study the transcriptional and translational effects of widespread pseudogene accumulation.

## RESULTS

### *Symbiopectobacterium endo*. maintains a greater load of younger, more recently formed, pseudogenes

The genomes of *Sodalis endo*. and *Symbiopectobacterium endo*. are 3.7 Mb and 4.5 Mb, respectively (Garber et al., 2021). Both genomes have many pseudogenes and appear to be in the early stages of genome erosion (**Table 1**). *Sodalis endo*. is closely related to the free-living bacteria *Sodalis praecaptivus* HS1 (Clayton et al., 2012; Chari et al., 2015), with which it shares about 90% genome-wide average nucleotide identity (ANI). Assuming an ancestral genome size similar to *S. praecaptivus*, the symbiont *Sodalis endo*. has lost about 30% of its genome. *Symbiopectobacterium endo*. has no close genus-level relatives that are free-living, but is phylogenetically within the *Brenneria*/*Pectobacterium* clade. The *Symbiopectobacterium* genome is close in length to the average 5 Mb-sized genomes seen across this clade, suggesting that it may have recently diverged from a free-living bacterium.

**Table 1:** Summary of pseudogenes and coding densities across three bacterial genomes studied here. The numbers in parentheses for the Intact genes and pseudogenes columns represent the amount of genome taken up by these annotations. Mb = megabase pairs, Kb = kilobase pairs. *Intergenic DNA excluding pseudogenes. **Coding density calculations exclude non-protein-coding genes.

| Bacterium | Genome size | Intact protein-coding genes | Pseudogenes | Non-coding DNA* | Coding density** |
| --- | --- | --- | --- | --- | --- |
| *Sodalis praecaptivus* | 5.159 Mb | 4331 (4.1 Mb) | 117 (130 Kb) | 929 Kb | 79.5% |
| *Symbiopectobacterium endolongispinus* | 4.492 Mb | 2559 (2.0 Mb) | 2679 (1.8 Mb) | 678 Kb | 44.5% |
| *Sodalis endolongispinus* | 3.728 Mb | 2294 (1.9 Mb) | 1807 (1.1 Mb) | 730 Kb | 51.0 % |

While both symbionts have similarly low coding densities of ∼45-50%, they differ in the types of pseudogenes they carry on their genomes (**Table 2**). We divided symbiont pseudogenes into three broad (and sometimes overlapping) categories: near-complete, truncated, and cryptic. Near-complete pseudogenes are very similar in size and structure to homologous functional genes on other genomes, but contain one or more indels or in-frame stop codons that disrupt the ancestral reading frame into two or more open reading frames (ORFs). Truncated pseudogenes are those where less than 65% of the ancestral gene length remains on the genome. Cryptic pseudogenes are those that retain few features of a functional gene and are difficult to detect by means other than sensitive ORF-independent sequence alignment algorithms. In both symbionts, the most common form of pseudogene is of the truncated type (**Table 2**).

**Table 2:** Broad categories assigned to pseudogenes in young endosymbionts from *P. longispinus*. The most numerous pseudogenes in both endosymbionts and *S. praecaptivus* are of the truncated type, and about 19% of pseudogenes are cryptic in the two symbionts *Symbiopectobacterium* and *Sodalis*. Near-complete pseudogenes, generally considered as the youngest (most recently formed) pseudogenes comprise a third (33%) of all pseudogenes in *Symbiopectobacterium*, but only 16% in *Sodalis endo*. and 4% in *S. praecaptivus*.

| Pseudogene category | Number in *Symbiopectobacterium* | Number in *Sodalis endo.* | Number in *S. praecaptivus* | Number in *Tremblaya princeps* |
| --- | --- | --- | --- | --- |
| *Cryptic* | 499 (19%) | 331 (19%) | 27 (23%) | 10 (53%) |
| *Truncated* | 1285 (48%) | 1173 (65%) | 82 (70%) | 5 (26%) |
| *Near-complete* | 895 (33%) | 303 (16%) | 8 (7%) | 4 (21%) |

We assume that small sequence changes—a small frameshift-inducing insertion or deletion, or a single-base substitution that changes a sense codon to a nonsense codon—occur more frequently than larger ones, and are thus more likely to initially break a gene (Liu et al., 2004; Kuo and Ochman, 2010; Danneels et al., 2018). As newly broken genes accumulate more deletions over time, they eventually appear as partial remnants of genes with major portions of the original ORF missing or unrecognizable because of lack of purifying selection to maintain the coding sequence. Following this assumption, we find that in comparison with *Sodalis endo*., *Symbiopectobacterium endo*. has a greater density of the (assumed) younger near-complete pseudogenes (**Table 2**). While this suggests that *Sodalis endo*. is an older endosymbiont, other mechanistic factors may be at play, such as the presence of numerous active transposases on the *Sodalis endo*. genome (Garber et al., 2021), which can catalyze deletions, including partial gene deletions (Plague et al., 2008), resulting in faster clearance of near-complete pseudogenes. Nevertheless, overall, it appears that *Symbiopectobacterium endo*. has a larger fraction of the more recently formed, or younger, class of near-complete pseudogenes.

To understand the downstream molecular consequences of a genome encoding large numbers of pseudogenes, we sequenced whole transcriptomes from mealybug bacteriomes (the bacteria-housing organs) to measure transcription across these endosymbiont genomes.

### Pseudogenes are widely transcribed and largely retain their ancestral transcription level

RNA sequencing reveals that endosymbiont pseudogenes continue to be transcribed, but, on average, at lower levels than intact genes. Out of 2679 pseudogenes encoded on the *Symbiopectobacterium endo*. genome, we detect 1978 (74%) within its transcriptome. We detect transcription of 2278 of 2559 (89%) of intact genes from *Symbiopectobacterium endo*. However, despite comparable numbers of intact genes (2278) and pseudogenes (1978) in the transcriptome of *Symbiopectobacterium endo*., pseudogenes are expressed at significantly lower levels compared to intact genes (**Figure 1A**) making up only 28% of the transcriptome by RNA read abundance, measured with normalized transcripts per million (TPM). This is also the case in *Sodalis endo*., where pseudogenes are not only expressed at lower levels (**Figure 1B**), but are also less likely to be transcribed. We detected 1152 of 1807 (64%) pseudogenes from *Sodalis endo*., which are also less abundant in the transcriptome, making up 19% by TPM-normalized read abundance.

**Figure 1:**
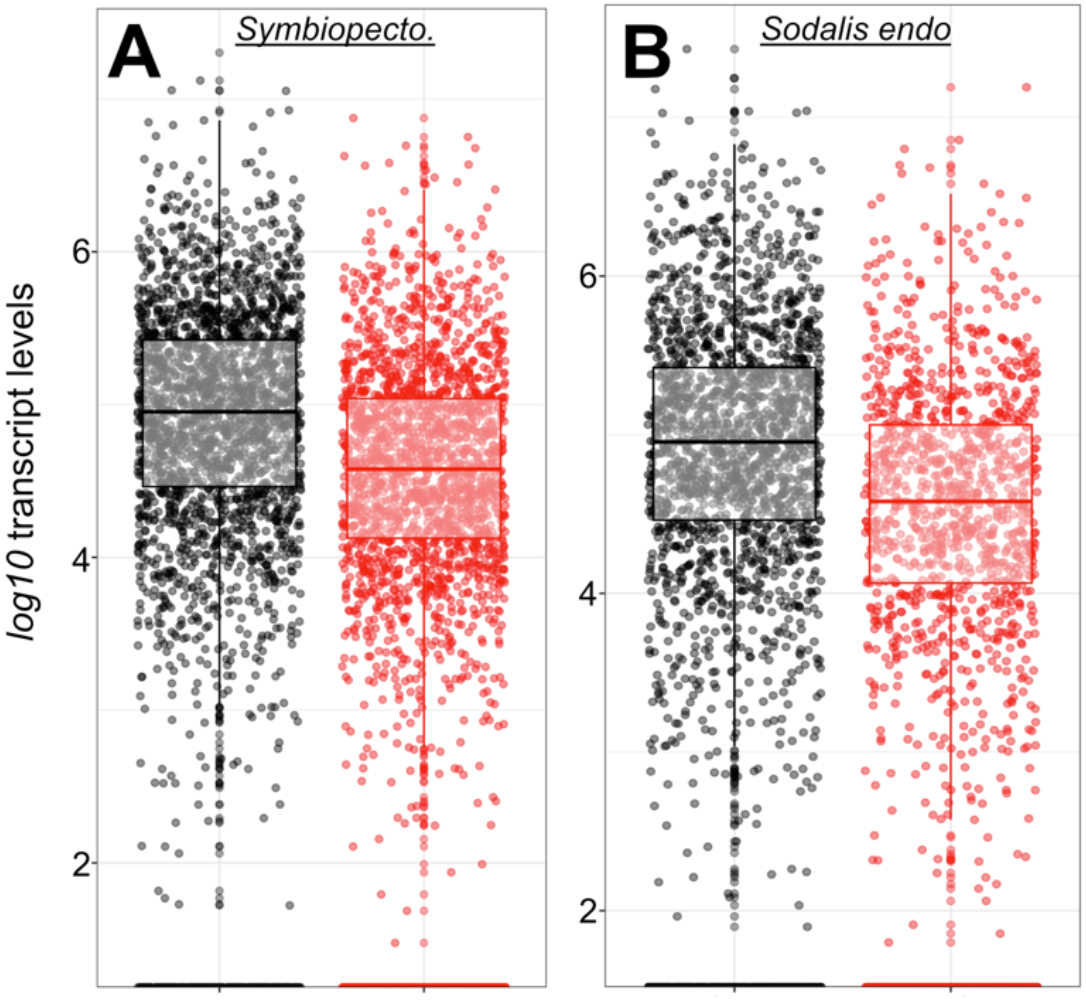
Transcript levels in A) *Symbiopectobacterium endo*., B) *Sodalis endo*. of intact genes (black dots) and pseudogenes (red dots). Welch’s t-test was applied to compare the expression profiles of intact genes and pseudogenes: p-values=8.69E-43 for *Symbiopectobacterium* and 7.57E-25 for *Sodalis endo*.

The lowered abundance of pseudogenes in both symbionts’ transcriptomes suggests that pseudogene transcripts may be inhibited or downregulated in some way, or, in pseudogenes that have been around longer, the RNA polymerase binding sites may have eroded away on the genome. However, it also could be that pseudogenes may simply be more likely to arise from lowly expressed genes. Given the close relationship between the free-living *Sodalis praecaptivus* and *Sodalis endo*., we had the unique opportunity to measure the expression levels of gene homologs that were pseudogenized on an endosymbiont genome but intact on a genome from a free-living close relative (**Figure 2A**). Of the 2128 orthologs shared between these two *Sodalis* genomes, 677 are pseudogenes in *Sodalis endo*. but remain intact in *S. praecaptivus*. In *S. praecaptivus*, these 677 genes are expressed at lower levels compared to genes that appear intact in *Sodalis endo*. (**Figure 2B**), supporting the hypothesis that lowly expressed genes are more likely to become pseudogenes. This pattern is consistent with previous work showing that highly expressed genes experience stronger purifying selection than lowly expressed genes (Yannai et al., 2018; Roberts and Josephs, 2023). It is also possible that highly expressed genes that become pseudogenes have a higher likelihood of causing negative impacts downstream, so are more quickly removed via selection (Kuo and Ochman, 2010). Consistent with the idea that lowly expressed genes are likely to become pseudogenes, we find that the (presumed younger) near-complete pseudogenes showed the lowest abundance levels of our three classes of pseudogene types compared with intact genes (**Supplemental Figure 1**). Overall, our data here suggest that what looks like lowered expression of pseudogenes compared to intact genes may simply reflect the fact that lowly expressed genes are more likely to become pseudogenes compared with highly expressed genes.

**Figure 2:**
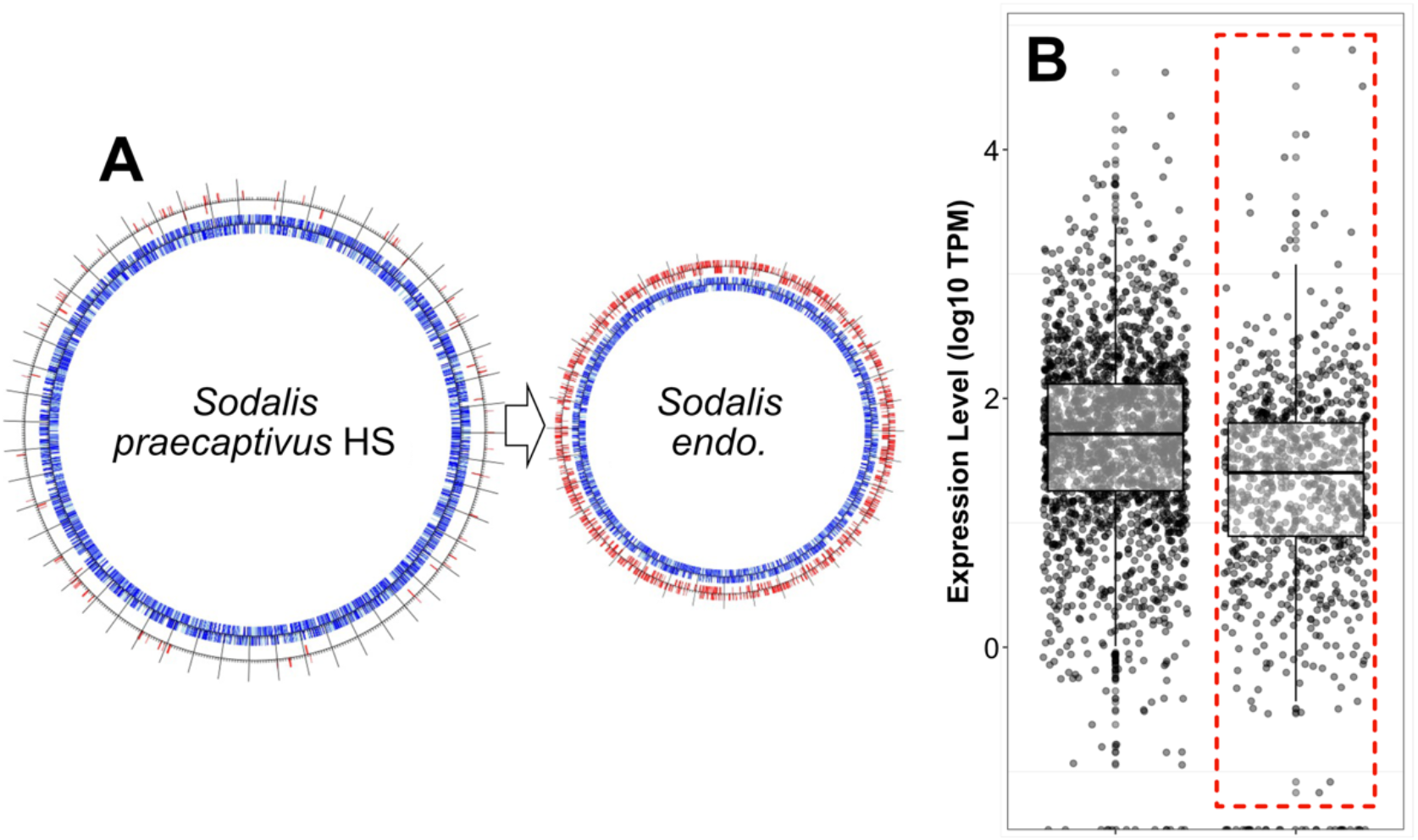
A) Circular genome diagrams of free-living *S. praecaptivus* and *Sodalis endo*. showing predicted gene regions as blue radial lines along the inner track. Pseudogenes are visualized on the outer track as red radial lines. B) Gene expression from *Sodalis praecaptivus*, showing a comparison between genes that remain intact and those that become pseudogenized on *Sodalis endo*, which are enclosed in a red-dotted box. Welch’s t-test was applied to test the differences between expression of to-become-pseudogenes and to-remain-intact genes in Panel B: p-value= 6.38E-32.

Having shown that a significant amount of pseudogene expression still occurs in these two endosymbionts, it seemed possible that these transcripts could still bind to ribosomes and attempt translation, with potentially deleterious downstream consequences. To explore this possibility, we next sequenced RNA transcripts that were bound to ribosomes using a modified ribosome profiling method for bacteria (Ingolia et al., 2009; Mohammad and Buskirk, 2019).

### Pseudogene transcripts continue to bind ribosomes in *Symbiopectobacterium endo*

Previous studies have found persistent transcription of pseudogenes combined with comparatively little protein products resulting from these broken genes (Goodhead et al., 2020; Nechitaylo et al., 2021; Feng et al., 2022). These patterns suggest that one or more post-transcriptional mechanisms are acting to prevent large amounts of non-functional and misfolded proteins from being produced. In search of possible mechanisms preventing the accumulation of large amounts of pseudogene-derived proteins, we isolated ribosomes from *P. longispinus* using sucrose gradient ultracentrifugation and sequenced all RNA that co-purifies with ribosome fractions. Our assumption here is that mRNA that co-purifies with ribosomes is likely to be actively translated, allowing us to infer the translational activities of *Sodalis endo*. and *Symbiopectobacterium endo*.

Ribosome-bound transcripts from *Symbiopectobacterium endo*. ribosomes dominated our sequencing efforts, yielding on average 30x more reads compared to *Sodalis endo*. (**Table 3**). The relative abundance of *Symbiopectobacterium endo*. RNA from ribosome-bound samples is inconsistent with previous measurements of the relative abundance of the two endosymbionts as assayed by genome coverage and RNA-FISH microscopy imaging (Garber et al., 2021), as well as the whole-transcriptome RNA sequencing and proteomic experiments performed in this study (**Table 3**). We attempted our total RNA isolations both with and without the translation inhibiting antibiotic chloramphenicol, and in both experiments obtained similar imbalances favoring *Symbiopectobacterium endo*. We suspect that the relative underabundance of ribosome-bound mRNA from *Sodalis endo*. is because *Sodalis endo*. ribosomes are less stable than *Symbiopectobacterium endo*. ribosomes in our purification protocol. Unfortunately, we do not know if this is an experimental artifact (*Sodalis endo*. ribosomes are less stable in the buffers we use) or whether it reflects actual biology (*Sodalis endo*. ribosomes are present but are less likely to be in active 70S conformations). While we report below on the RNAs bound to *Sodalis endo*. ribosomes, we caution that these results may be unreliable or biased in some way that we cannot predict.

**Table 3:**
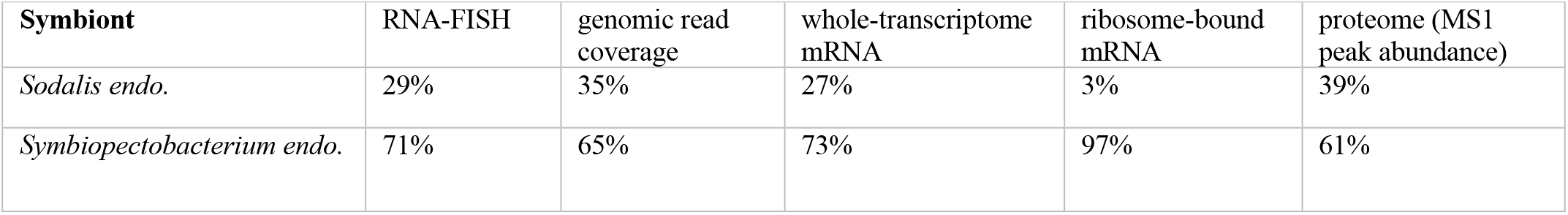
Relative abundances of various biomolecules in the two intra-*Tremblaya* endosymbionts. <u>RNA-FISH</u> is a proxy for SSU rRNA; <u>genomic read coverage</u> measures the relative number of genome copies, <u>whole-transcriptome mRNA</u> measures the total amount of transcripts, while ribosome-bound mRNA measures the number of ribosome-bound transcripts undergoing translation; proteome abundance is measured as a proxy of peptide peak abundance from the initial mass spectrometry scan (MS1).

| Symbiont | RNA-FISH | genomic read coverage | whole-transcriptome mRNA | ribosome-bound mRNA | proteome (MS1 peak abundance) |
| --- | --- | --- | --- | --- | --- |
| *Sodalis endo.* | 29% | 35% | 27% | 3% | 39% |
| *Symbiopectobacterium endo.* | 71% | 65% | 73% | 97% | 61% |

We find further contrast between the two symbionts in the proportions of pseudogene transcripts bound to their ribosomes. *Symbiopectobacterium* attempts translation of nearly all of its pseudogene transcripts, with similar counts of intact (2202) and pseudogene (1923) transcripts bound to ribosomes (orange dots in **Figure 3A**). By contrast, only 151 out of a total 786 detected pseudogene transcripts were found among the ribosome-bound RNA in *Sodalis endo*. (orange dots in **Figure 3B**). In both endosymbionts, ribosome-bound RNA from pseudogene transcripts is less abundant compared with intact gene transcripts (22% in *Symbiopectobacterium endo*. and 8% *Sodalis endo*.) (**Supplemental Figure 2**), suggesting that transcripts encoding pseudogenes are less likely to stably bind ribosomes or are more likely to terminate prematurely, or both.

**Figure 3:**
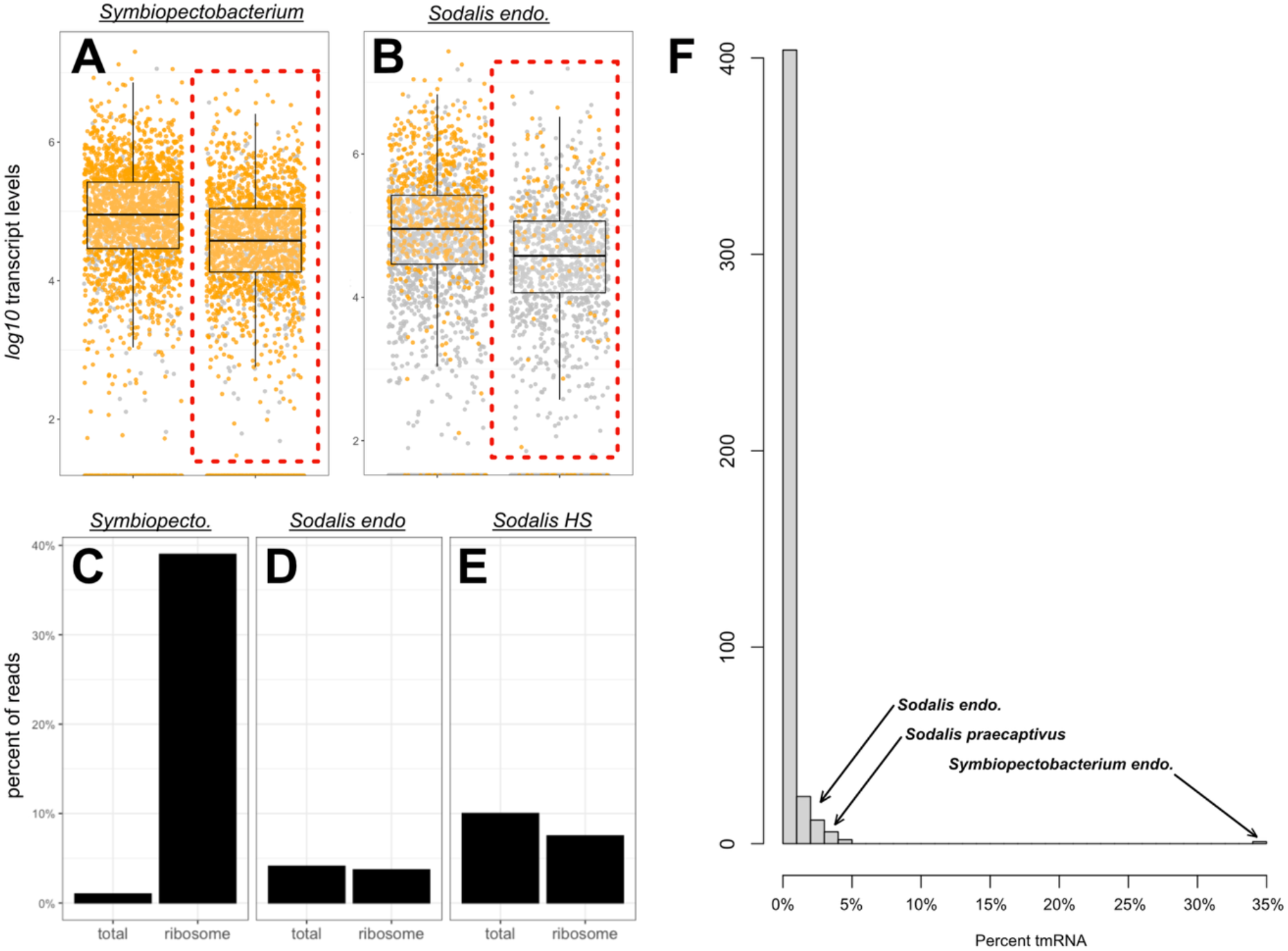
Gene expression levels in A) *Symbiopectobacterium endo*. and B) *Sodalis endo* (same data as in Figure 1). Pseudogenes are highlighted on the right side of each plot with red dotted boxes. Dots filled in with yellow indicate genes identified among the ribosome-bound RNA. Panels C-E show the relative amounts of tmRNA among the total and ribosome-bound transcriptomes in C) *Symbiopectobacterium endo*., D) *Sodalis endo*, and E) *Sodalis praecaptivus*. F) Histogram showing the percent of total ribosome-associated RNA that mapped to tmRNA from 449 sequencing studies, with arrows pointing to where three of our own data points fall.

Surprisingly, nearly 40% of all RNA obtained from *Symbiopectobacterium endo*. ribosomes is from the transfer-messenger RNA (tmRNA; **Figure 3C**). This small RNA, which contains both an mRNA and a tRNA-like domain (Keiler et al., 1996), is involved in recycling bacterial ribosomes that have stalled on broken mRNA transcripts (Moore and Sauer, 2005; Janssen and Hayes, 2012; Keiler, 2008). The tmRNA gene is conserved across bacteria, and transcripts from it are often the most abundant and stable RNAs in the cell (Ray and Apirion, 1979), though there are rare cases in which either the protein or RNA component of the tmRNA system has been lost in endosymbionts with highly reduced genomes (Hudson et al., 2014). In *Symbiopectobacterium endo*., tmRNA comprises about 1% of the total transcriptomic RNA, and so its presence as 40% of all ribosome-bound transcripts is significant. In both *Sodalis endo*. and the free-living *Sodalis praecaptivus*, the enrichment of tmRNA on ribosomes vs. total transcriptomes is less pronounced (3.7% vs. 4.1% in *Sodalis endo*. and 7.4% vs. 10% in *Sodalis praecaptivus*, **Figure 3D-E**). The overabundance of tmRNA among the ribosome-bound RNA fraction appears especially pronounced when we compared it to the tmRNA abundance in other comparable datasets from ribosome profiling and footprinting data (**Figure 3F, Supplemental Table 1**).

The large number of pseudogene transcripts binding to ribosomes, particularly in *Symbiopectobacterium endo*., raises the possibility that high levels of non-functional or toxic protein products could accumulate in these cells. To investigate whether or not proteins were produced from these ribosome-bound pseudogene transcripts, we performed shotgun mass spectrometry (MS) proteomics on mealybug bacteriomes.

### Protein products from pseudogenes are rare, particularly in *Symbiopectobacterium endo*

MS proteomics of mealybug bacteriomes yielded 6041 proteins from the host mealybug, which comprised the vast majority of our MS data. The rest of the proteome is split roughly evenly between *Tremblaya* (120 proteins, 6% of peptides by abundance), *Sodalis endo*. (597 proteins, 5% by abundance) and *Symbiopectobacterium endo*. (558 proteins, 4% by abundance). Among the 558 proteins detected from *Symbiopectobacterium endo*., only 42 are pseudogenes (**Table 4, Supplemental Table 2**), and all are of the truncated type. Based on the relative abundance of each identified peptide measured via intensity of MS1 precursor ions (Palomba et al., 2021), pseudogene-derived peptides are in low abundance, making up less than 3% of the total *Symbiopectobacterium endo*. proteome (**Figure 4A**). The majority of the *Symbiopectobacterium endo*. proteome consists of peptides derived from intact transcripts that are abundant among isolated ribosomes (**Figure 4B**). In *Sodalis endo*., by contrast, we find little relationship between ribosome-bound transcript levels and protein presence, and pseudogene-derived peptides, from 77 truncated pseudogenes, comprise 11% of the proteome by abundance (**Figure 4C-D, Supplemental Figure 3**).

**Table 4:**
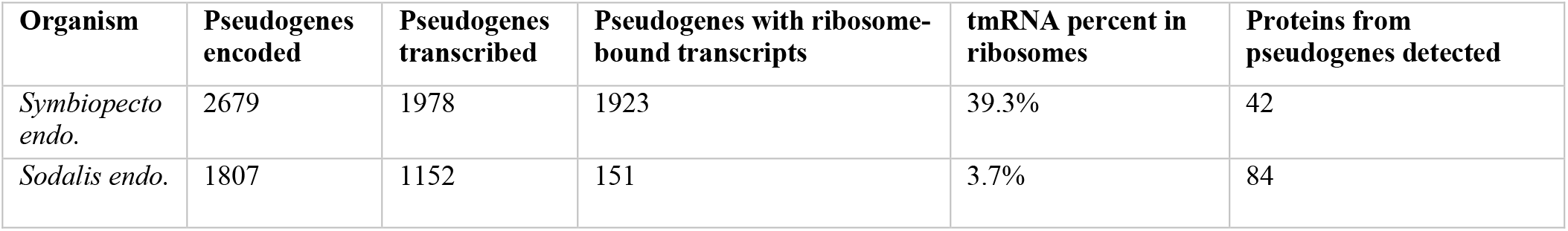
Summary of pseudogenes across the symbionts’ genomes, transcriptomes, and ribosome-bound RNA. tmRNA abundance is shown as a percentage of total mRNA translation (excluding tRNAs, rRNAs, and other non-coding RNA sequences).

**Figure 4:**
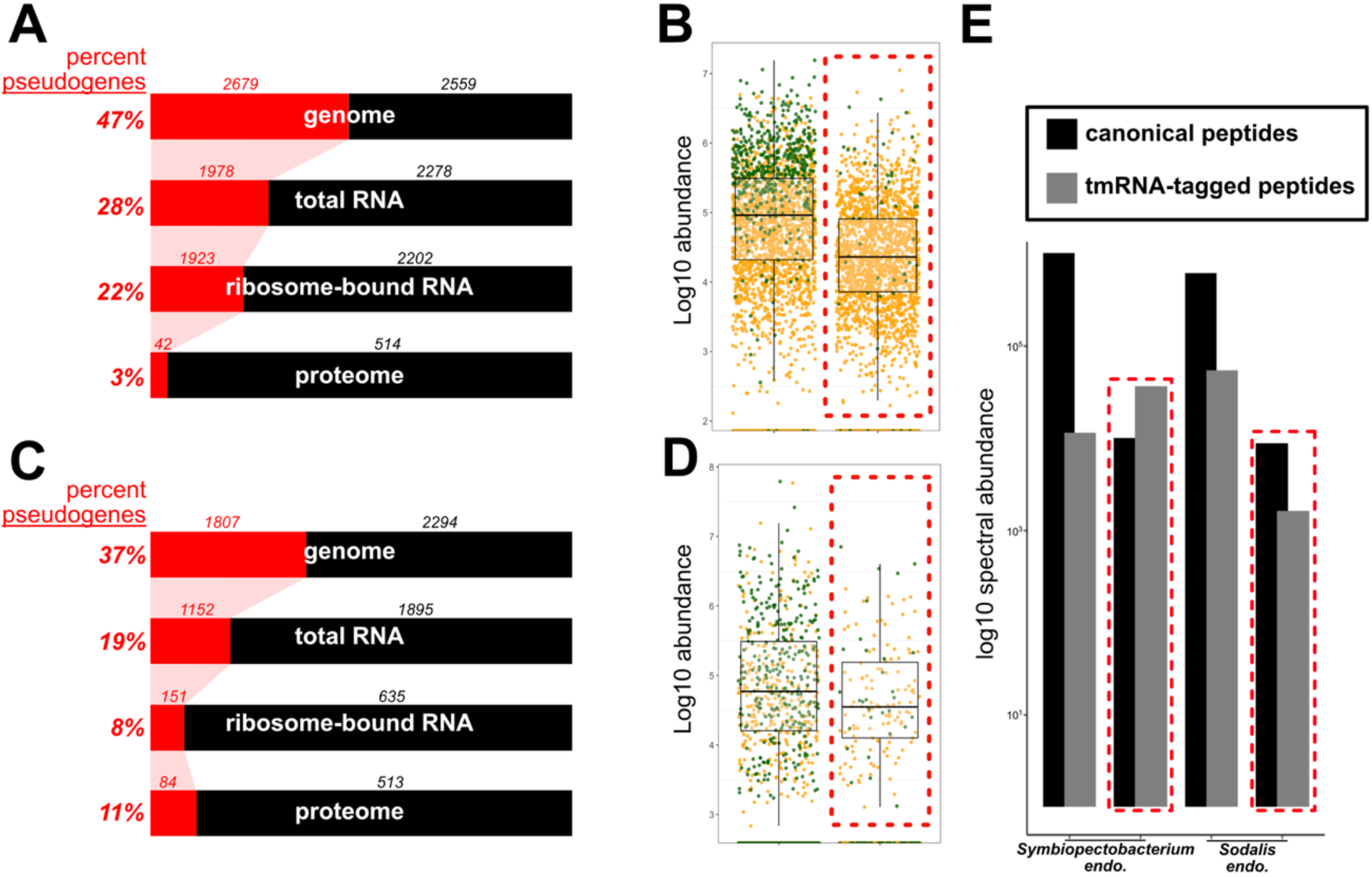
A) Sankey diagram showing the relative proportion of each biomolecule that corresponds to pseudogenes (shown in red) in Symbiopectobacterium. For genome, percent pseudogene refers to proportion of bases covered by pseudogenes. For the total and ribosome-bound RNA, percent pseudogene corresponds to percent of total RNA abundance. For proteome, percent pseudogene corresponds to percent of total peptides by abundance. Small numbers above each Sankey bar indicate the numbers of genes or pseudogenes identified at each stage. B) Translation levels in *Symbiopectobacterium endo*., with pseudogenes denoted in red-dotted boxes and transcripts whose peptides are detected in the proteome colored green. C) Sankey diagram for Sodalis endo. D) Translation levels in *Sodalis endo*., with pseudogenes in red-dotted boxes and transcripts whose peptides are detected in the proteome colored green. E) Relative peptide abundance of canonical and tmRNA-tagged peptides in *Sodalis* and *Symbiopectobacterium endo*. Spectral abundance values are shown on a log-scale, with arbitrary spectral units obtained from the mass spectrometer. Pseudogenes denoted in red-dotted boxes. Welch’s t-test was applied to ribosome-bound RNA levels of intact genes vs pseudogenes: p-value=2.90E-71 for *Symbiopectobacterium* and 0.68805323 for *Sodalis endo*.

Given the high abundance of tmRNA bound to *Symbiopectobacterium endo*. ribosomes, we looked for potential products of ribosome rescue in the proteome. The coding region of tmRNA codes for a short amino acid tag that is appended to proteins stalled on broken transcripts, targeting them for degradation (Keiler, 2008). When we included peptides with this tag in our proteomic search space, we identified additional peptides bearing the ANDENYALAA tag in *Symbiopectobacterium endo*. and the ANDSQFESKTALAA tag in *Sodalis endo*. These tmRNA-tagged peptides are in relatively low abundance compared to unmodified peptides, except from *Symbiopectobacterium endo*. pseudogenes (**Figure 4E**). Based on relative spectral abundances, pseudogene-derived peptides with the tmRNA tag outnumber unmodified pseudogene peptides 4-fold in *Symbiopectobacterium endo*. Overall, these proteomic data suggest a higher level of tmRNA activity in *Symbiopectobacterium endo*., consistent with the large amount of tmRNA that we find co-purifying with its ribosomes (**Figure 3C**).

## DISCUSSION

### Short-term vs. long-term adaptations to large numbers of pseudogenes

When a pseudogene is first formed, the upstream sequence elements involved in RNA polymerase and ribosome recognition are unchanged, and therefore transcription and translation should attempt to proceed as normal. Depending on the nature of the brand-new pseudogene, the fitness effect of continued transcription and translation might vary from minor, such as a deletion that affects the very end of a lowly expressed gene, to major, such as a deletion that alters the reading frame near the middle of a highly expressed gene. Over evolutionary time, selection will eliminate the sequence elements of this pseudogene that allow it to be transcribed and translated. We see evidence of this type of evolution in the global analysis of start codons and Shine-Dalgarno (ribosome-binding) sites upstream of pseudogenes compared to functional genes. We find that pseudogenes are less likely to have upstream Shine-Dalgarno sites (**Supplemental Figure 4**) and are also less likely to start with the preferred ATG initiation codon (Belinky et al., 2017) (**Supplemental Figure 5**). Of special interest to us in this study was to uncover any short-term mechanistic responses that bacteria might use to repress transcription and translation of pseudogenes over short time frames, before evolution has had time to eliminate the RNA polymerase and ribosome binding sites. For this reason, we were especially interested in the mechanisms that *Symbiopectobacterium endo*. might use to prevent the formation of aberrant proteins, because our analysis of pseudogene age suggests that this endosymbiont carries a larger fraction of newer pseudogenes on its genome (**Table 2**).

### Lowly expressed genes are more likely to become pseudogenes

We reasoned that there might not be a strong mechanistic response to pseudogene formation at the level of transcription. While some mechanisms are known to affect mRNA stability in bacteria (Mohanty and Kushner, 2016), we know of no mechanism in bacteria similar to nonsense-mediated decay in eukaryotes (Chang et al., 2007) that might alter the stability of pseudogene transcripts specifically, although the tmRNA system has been shown to be directly involved in degrading aberrant mRNAs (Yamamoto et al., 2003 and Venkataraman et al., 2014). When looking at the relative abundance of transcripts from functional genes and pseudogenes globally, we, like others before us (Goodhead et al., 2020; Nechitaylo et al., 2021; Feng et al., 2022), see an average decrease in the levels of pseudogenes relative to transcripts from intact protein-coding genes (**Figure 1A-B**). Some of this reduced expression is likely a consequence of the loss of transcription binding sites due to sequence evolution of older pseudogenes. However, we wondered how much of this signal was simply due to the higher probability of lowly expressed genes becoming pseudogenes, rather than some global mechanistic response that would reduce pseudogene expression. The idea here is that, on average, lowly transcribed genes are generally under weaker purifying selection (Yannai et al., 2018; Roberts and Josephs, 2023) and therefore, might be less likely to be important in the context of endosymbiosis. To test this hypothesis, we compared transcript abundance of *Sodalis endo*. pseudogenes to their intact orthologs from *Sodalis praecaptivus*. We found that genes destined to become pseudogenes in *Sodalis endo*. are lower in abundance within the transcriptome of *Sodalis praecaptivus* (**Figure 2B**).

Along with the fact that no known mechanism for systematically down-regulating pseudogene transcripts in bacteria exists, these results suggest that there is little to no transcriptional response to becoming a pseudogene on short time scales: transcription continues as normal until sequence evolution alters transcriptional binding sites. Weaker transcription of pseudogenes in *Sodalis endo*. compared to *Symbiopectobacterium endo*. (**Table 4**) may thus reflect a difference in how long each symbiont has been evolving under weakened selection, with *Symbiopectobacterium endo*. representing the younger endosymbiont, captured before most of the degradation of pseudogene transcriptional signals has had time to occur.

### Transcripts from pseudogenes bind ribosomes but rarely make protein products

Before sequence evolution has time to change the promoter sequences of a pseudogene, we expect it to be transcribed as usual and for the transcripts to bind ribosomes as usual. We see evidence of this in our purified ribosome preparations, where a substantial amount of *Symbiopectobacterium endo*. pseudogene transcripts are found bound to ribosomes (**Figure 3A**). However, very few of those transcripts are made into protein products, or very few of these protein products remain stable or abundant enough in the cell for us to measure in mass spectrometry experiments (**Figure 4A**). What mechanism might be at work to prevent pseudogene transcripts from being made into aberrant protein products, before evolution has had time to erode away the ribosome binding signals? The high amounts of tmRNA from *Symbiopectobacterium endo*. in our purified ribosome preparations suggest a possible mechanism. The tmRNA is known to be abundant in bacterial RNA-seq datasets (Engelhardt et al., 2020), but we were unsure how abundant it was in ribosome profiling experiments. We analyzed ribosome profiling data from *Escherichia coli* (Mangano et al., 2022), as well as *Streptomyces venezuelae* and *Streptomyces griseus* (Kim et al., 2020), which were generated using methods similar to ours (direct sequencing of ribosome-bound RNA without micrococcal nuclease digestion). The percent of reads mapping to tmRNA ranged from 0.02% in *S. griseus*, to 0.44% in *S. venezuelae*, to 3.4% in *E. coli* (**Supplemental Table 1**). Other studies using ribosome footprinting showed similar proportions of tmRNA, never exceeding 4.7% of total ribosome-protected RNA fragments (**Figure 3F**). These data suggest that our measurement of over 39% of reads from *Symbiopectobacterium* being from tmRNA is significant (**Figure 3C, Table 4**), though we cannot fully rule out artifacts introduced from differences in protocols, or the fact that our system is considerably more complex than bacterial isolate cultures.

The function of tmRNA is to rescue stalled ribosomes that have attempted translation on a broken mRNA transcript. The tmRNA first enters the ribosomal aminoacyl-tRNA binding site via its tRNA-like domain, causing the ribosome to switch over to the templated mRNA sequence of tmRNA and to translate a short 10-amino acid tag that is appended to the nascent protein (Moore and Sauer, 2005). This mechanism is colloquially known as ribosome rescue and is thought to allow bacteria to quickly recycle inefficient or stalled ribosomes (Moore and Sauer, 2005). We found evidence for products of active ribosome rescue (protein fragments appended with a short amino acid tag encoded on the tmRNA CDS region) in our mass spectrometry data, in particular in *Symbiopectobacterium endo*., where there are about 4-fold more pseudogene peptides with the tmRNA tag than without (comprising 78% of pseudogene-derived peptides). This is in contrast to products of *Symbiopectobacterium* intact genes, where only one percent of peptides are found with a tmRNA tag. By contrast, in *Sodalis endo*., rescue-derived products are in the minority among peptides from both intact genes and pseudogenes, but still enriched among the pseudogene products.

### Proposed mechanism for silencing of newly formed pseudogenes

Our finding that large amounts of tmRNA are bound to *Symbiopectobacterium endo*. ribosomes suggests a role for this RNA in eliminating aberrant pseudogene-derived proteins from the proteome. However, tmRNA only acts on broken mRNA transcripts, and so at least one more step is required for our model to work. It has been shown that ribosome collisions can recruit the activity of tmRNA due to the cleavage of mRNA on collided ribosomes upstream of the stall site by the protein SmrB, which then allows tmRNA and its partner protein SmpB to enter the ribosome and terminate translation (Saito et al., 2022). Both the SmrB and SmpB proteins and their transcripts were detected in our mass spectrometry and transcriptomic data (**Supplemental Figure 6**). It seems reasonable to hypothesize that pseudogenes might experience ribosome collisions more frequently, due to shifts in codon bias that occur when an open reading frame is perturbed. This subtle consequence of codon usage changes, which can be observed when we compare the codon usage of pseudogenes in *Sodalis endo*. with their intact orthologs in *S. praecaptivus* (**Supplemental Figure 7**), may have a significant impact on translation efficiency and stalling due to higher incidences of rare codons, premature stop codons (**Supplemental Figure 8**), and cognate tRNA scarcity (Roche and Sauer, 1999; Samatova et al., 2020). We emphasize that we do not yet have direct experimental evidence that pseudogenes are more likely to result in ribosome collisions and stalling, but this seems like an interesting path for future research.

Overall, our data suggest that bacterial symbionts do little to prevent the transcription and translation of pseudogenes as they first emerge. Over time, sequence evolution away from preferred RNA polymerase and ribosome binding sites will decrease or prevent the transcription and translation of pseudogenes. How bacteria prevent newly formed pseudogenes from producing products has been less clear. Here, we present data suggesting the ribosome rescue system as a mechanism to both rapidly degrade proteins made from pseudogenes and rescue ribosomes that are pausing, stalling, and colliding on non-optimal sets of codons. This ribosome rescue system may be useful for bacteria that have found themselves in a state where many pseudogenes exist on their genome but the time needed to eliminate ribosomal initiation sites through sequence evolution has not yet passed.

## MATERIALS AND METHODS

### Insect rearing and bacterial culturing

*Pseudococcus longispinus* populations were reared on sprouted potatoes at 25°C, 77% relative humidity, and a 12 h light/dark cycle in a Percival 136LL incubator. *Sodalis praecaptivus* HS1 was grown on liquid lysogeny broth (LB, Miller) at 30 °C with vigorous shaking.

### RNA extraction and sequencing of mealybugs and *Sodalis praecaptivus*

For transcriptomic sequencing, *P. longispinus* mealybug bacteriomes were dissected, immediately flash frozen in liquid nitrogen, and then crushed with a plastic pestle inside sterile microcentrifuge tubes (1.5 mL). Tissue lysates were clarified (to remove insoluble cell debris) with a 30 min spin at 20,000 g (4°C). RNA from clarified lysates was purified using the Qiagen RNeasy kit with 1 mM DTT added to the lysis buffer (Buffer RLT), along with 0.0002 U/µl RNase Inhibitor (Thermo Fisher Scientific). We performed DNA digestion with DNase I prior to washing and elution with double-distilled H2O (ddH_2_O). Purified RNA quality was assessed using the Agilent 4200 TapeStation. Depletion of ribosomal RNA was performed with Illumina’s RiboZero kit using standard probes. Sequencing was performed by Genewiz with the Illumina NextSeq 500 platform.

*Sodalis praecaptivus* HS cells were harvested at mid-log phase of growth (OD600 between 0.6 and 0.9) with 15 min of centrifugation at 5000 g (4°C). Cells were lysed by flash-freezing in liquid nitrogen and passing through a 27-gauge syringe 15-20 times. Lysates were clarified as discussed above and RNA extracted following the same protocol as mealybug samples.

### Ribosome profiling of mealybugs and *Sodalis praecaptivus*

We performed ribosome profiling on whole mealybugs, as well as on dissected bacteriome tissue. In both cases we used ice-cold Buffer A (20 mM Tris-HCl pH 7.5, 100 mM NH_4_Cl, 10.5 mM Mg(OAc)_2_, 0.5 mM EDTA, 77 µM chloramphenicol), containing 0.0002 U/µl RNase Inhibitor, 0.0002 U/µl DNase I, and two tablets of Pierce Protease Inhibitor (Thermo Fisher Scientific) for lysis. Additionally, in the bacteriome preparation, we included 25 μg/mL chloramphenicol. Samples were homogenized using a Dounce homogenizer with 15-20 mechanical strokes. The resulting lysates were centrifuged at 20,000 g for 30 minutes (4°C). Pellets were discarded and supernatant (1 ml each) layered onto 12 mL 10-50% sucrose gradients made with Buffer B (20 mM Tris-HCl pH 7.5, 500 mM NH4Cl, 10.5 mM MgOAC, 0.5 mM EDTA, 1.1 M sucrose). We also added 2-mercaptoethanol (420 µl/L), benzamidine (1 ml/L of 100 mM), and phenylmethylsulfonyl fluoride (PMSF, 2 mL/L 50 mM) to both Buffer A and B immediately before use. The gradient was prepared with a BioComp Gradient Master, and tubes inserted into the Swinging Bucket 40 Ti rotor and spun for 2.5 h at 200,000 g, 4°C. Resulting gradients were cut using the BioComp Piston Gradient Fractionator, with elution absorbances measured at 260 nm to detect nucleic acids. Ribosomal fractions corresponding to 70S ribosomes and polysomes were taken for RNA extraction and sequencing (**Supplemental Figure 9**).

*S. praecaptivus* HS1 colonies were picked from LB agar plates and sub-cultured into conical flasks containing LB. Cells were harvested in the mid-log phase of growth and pelleted with centrifugation at 5000 g for 15 min (4°C) and homogenized using a high-pressure homogenizer (5 rounds, 20,000 psi). Homogenization was performed using Buffer A, with 0.0002 U/µl RNase Inhibitor, 0.0002 U/µl DNase I, two tablets of Pierce Protease Inhibitor, and 25 μg/mL chloramphenicol. The resulting lysates were clarified and subjected to sucrose-gradient ultracentrifugation as described above for mealybug samples. Elution absorbances measured at 260 nm yielded monosome peaks starting at fraction 10 (roughly halfway down in the gradient), and more clearly-defined polysome peaks in fractions 14-20 (**Supplemental Figure 10**).

RNA was extracted using TRIzol (Thermo Fisher Scientific) with 20% chloroform. Each fraction from the sucrose gradient (600 μl) was combined with 2.4 mL of TRIzol reagent, vortexed, and incubated for 5 minutes. Each fraction then received 480 μl of chloroform (Thermo Fisher Scientific), incubated for 3 minutes, and centrifuged for 15 min at 20,000 g (4°C). After centrifugation, the clear aqueous phase was placed into a sterile 15 mL conical tube and combined with 1.2 mL of 100% isopropyl alcohol, followed by overnight incubation at -20°C. Samples were then centrifuged at 20,000 g for 30 min (4°C), pellets washed twice with 75% ethanol, and resuspended in DNase/RNase free ddH_2_O. RNA quality was assessed with the Agilent 4200 TapeStation before being sent to SeqCoast Genomics for sequencing using the Illumina NextSeq 1000 platform. Ribosome depletion was performed using custom rRNA probes designed specifically for ribosomal RNA from *P. longispinus*, including the host and its symbionts (**Supplemental Table 3**). Standard probes were used for RNA from *Sodalis praecaptivus* HS1.

### Mass spectrometry proteomics of mealybugs

*P. longispinus* mealybug bacteriomes were dissected into phosphate-buffered saline and immediately flash-frozen in liquid nitrogen. Samples were processed at the Translational Genomics Research Institute (TGen) via biopulverization, followed by lysis in a Precellys homogenizer with soft tissue beads. Samples (approximately 50 μg of protein) were then subjected to in-solution trypsin digestion. Digested peptides were purified using solid-phase extraction on C18 columns dried in a SpeedVac. Peptides were then resuspended in 2% acetonitrile, 0.1% formic acid solution. For peptide fractionation, a kit-based high-pH reversed-phase method was used, yielding eight fractions. Peptides were separated over a 120-minute gradient on a 25 cm Aurora Ultimate C18 column (IonOpticks) using a Vanquish Neo UHPLC system coupled to a Thermo Scientific Orbitrap Eclipse mass spectrometer. Data-dependent acquisition was performed with scan range set to 375 - 1500 m/z. MS1 scans were acquired in the Orbitrap at a resolution of 120,000 FWHM (full width at half-maximum) and MS2 scans were collected in the ion trap.

### Bioinformatics

#### Pseudogene annotation

Pseudogenes were annotated with Pseudofinder v1.1.0 (Syberg-Olsen et al., 2022), using NCBI’s non-redundant protein database for reference. In cases where >65% of the gene is retained on the genome, but is broken up into multiple ORFs, that gets labeled as a near-complete pseudogene; we consider these near-complete pseudogenes to be the youngest, most recently formed. If 65% or less of a gene is present on the genome, determined in comparison to the average length of the top 50 orthologs from RefSeq, that gene gets labeled as a truncated pseudogene; we consider these pseudogenes to be generally older than near-complete pseudogenes. Finally, regions considered to be intergenic noise by the original gene-calling software used here (Prodigal v2.6.3 [Hyatt, 2010], via Prokka v1.14.5 [Seemann, 2014]) were aligned against the RefSeq database, and regions with at least 25 significant matches within that database are labeled as cryptic pseudogenes. We consider these pseudogenes to be older than truncated pseudogenes.

#### Transcriptomics and ribosomal profiling

RNA reads were quality-trimmed using Trimmomatic v0.39 (Bolger et al., 2014) and aligned against *Sodalis praecaptivus* and endosymbiont genomes using Bowtie2 v2.4.5 (Langmead and Salzberg 2012). Per-gene counts data were extracted using HTSeq-Count v0.11.2 (Anders et al., 2015) and count values were converted to normalized expression levels (correcting for differences in gene lengths, sequencing effort per sample, and symbiont abundance), expressed as transcripts per million (TPM, Zhao et al., 2021).

For estimation of reads from tmRNA in ribosome profiling datasets, we searched for similar datasets in NCBI’s Sequence Read Archive (SRA). We identified 40 BioProject studies with a total of 438 SRA experiments corresponding to RNA sequencing of ribosome and polysome isolations. Of these, two studies generated data using methods similar to ours (denoted with asterisks in **Supplemental Table 1**). We used prefetch v3.0.0 and fastq-dump v3.0.0 from the SRA Toolkit to download the datasets, and tmRNA reads were classified by alignment, using Bowtie2 v2.4.5 (Langmead and Salzberg, 2012), against an expanded database of tmRNA sequences from Bacteria and Archaea (Nawrocki et al., 2025). This was repeated for our own ribosome profiling dataset, where we found that 35% of the *Symbiopectobacterium*-classified reads mapped to the tmRNA database. This is only marginally lower than our estimates based on alignment of reads against the annotated *Symbiopectobacterium* tmRNA sequence.

#### Statistical comparisons and data visualization

We compared expression levels between pseudogenes and intact genes using Welch’s t-test (Welch, 1947), which allows for comparisons between groups of varying sizes (there are more intact genes than pseudogenes) and variances, though there does not appear to be much difference in expression level variation between intact and broken genes. This test was carried out in RStudio (Posit team, 2025), and we used a combination of base R and ggplot2 for visualization.

#### LC-MS/MS Data Processing

Raw data were processed with Proteome Discoverer (Thermo Fisher Scientific, v3.1). Spectra were searched with the Sequest HT search engine using the species-specific whole proteome FASTA database (enzymatic cleavage set to trypsin, allowing up to two missed cleavages). Precursor mass tolerance was set to 10 ppm and fragment mass tolerance was set to 0.6 Da. Peptide validation was carried out via the Percolator node (Spivak, 2009), which uses a machine-learning model trained on true (target) and false (decoy) identifications. We set the false discovery rate (FDR) to 1% (Elias and Gygi, 2007). For quantification, precursor-based feature detection and mapping were performed, and ion intensities were extracted for peptide-level abundance quantification. To detect products of ribosome rescue, we generated a custom protein database that comprised all possible combinations of peptide + tag from the tmRNA. While this resulted in a large protein database search space for Proteome Discoverer (since we included the tmRNA tag at all possible positions where a protein may stall), we maintained the 1% FDR cutoff to mitigate false positives.

All software versions are listed in **Supplemental Table 4**.

## Data Availability

Genomic data from *Pseudococcus longispinus* are available via BioProject PRJEB12068 (published in Garber et al., 2021). RNA sequencing data generated in this study have been made available via NCBI’s Sequence Read Archive (BioProject PRJNA1417914 for *Sodalis praecaptivus* RNA sequencing data and PRJNA1418813 for *P. longispinus* RNA data). Mass spectrometry data have been made available via the PRIDE Archive (PXD074041).

## Acknowledgements

We would like to thank Tim Karr of the ASU Mass Spectrometry Facility for helpful feedback on the analysis of mass spectrometry data and Rachel Green, Scott Miller, and Tim Wheeler for advice on design of the ribosome profiling experiment. This work was supported by the Howard Hughes Medical Institute and NASA’s Interdisciplinary Consortia for Astrobiology Research (grant number 80NSSC23K1357).

## SUPPLEMENTAL FIGURES

**Supplemental Figure 1:**
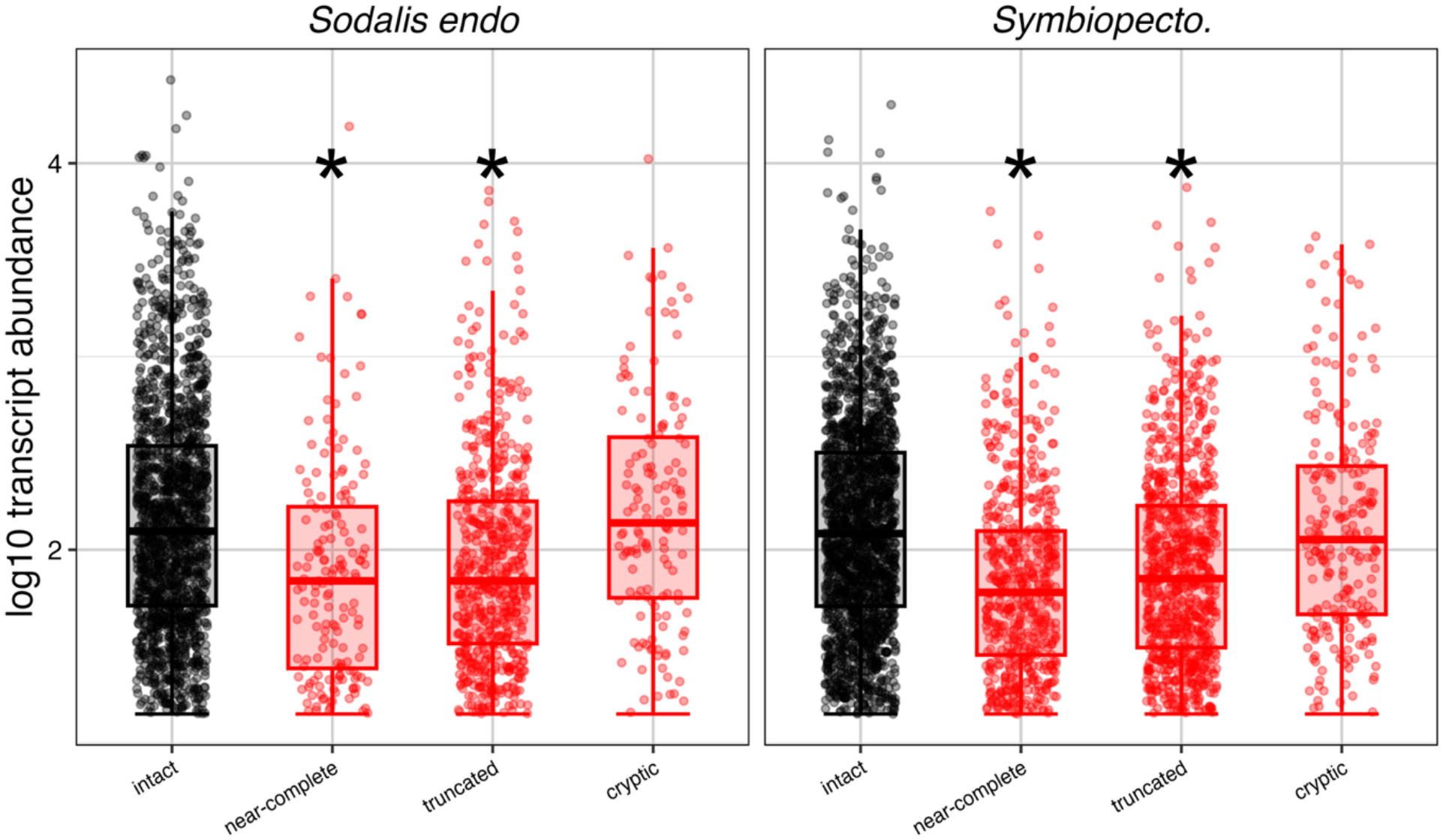
Transcript levels in *Sodalis endo* and *Symbiopectobacterium endo* of intact genes (black dots) and the different categories of pseudogenes (red dots). Welch’s t-test was applied to compare the expression profiles of the different types of pseudogenes with those of intact genes. Asterisks indicate where significant differences were predicted (p-value < 1.35E-06).

**Supplemental Figure 2:**
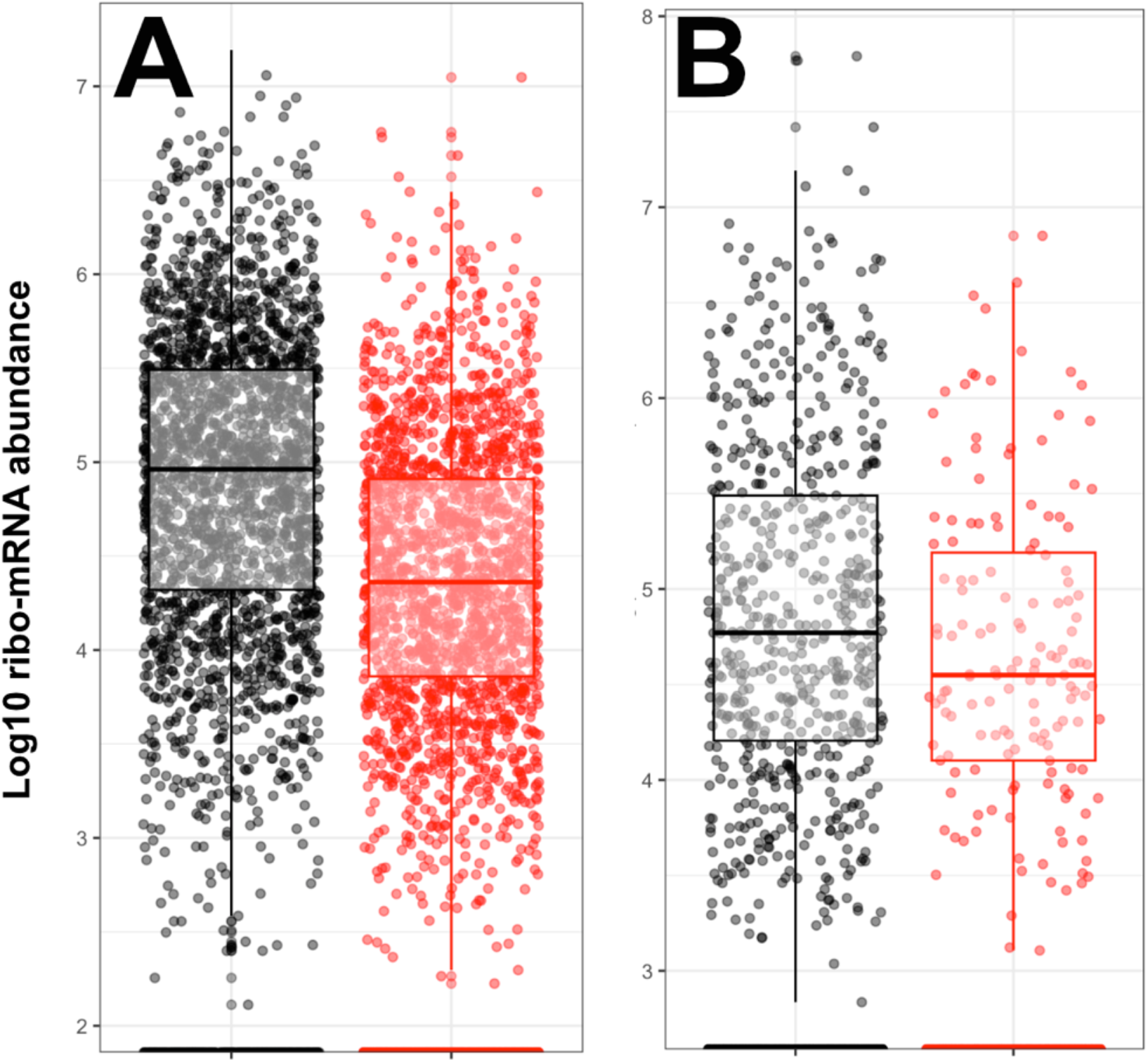
Normalized RNA levels among the ribosomes in A) *Symbiopectobacterium endo*. and B) *Sodalis endo*. Pseudogenes are colored red.

**Supplemental Figure 3:**
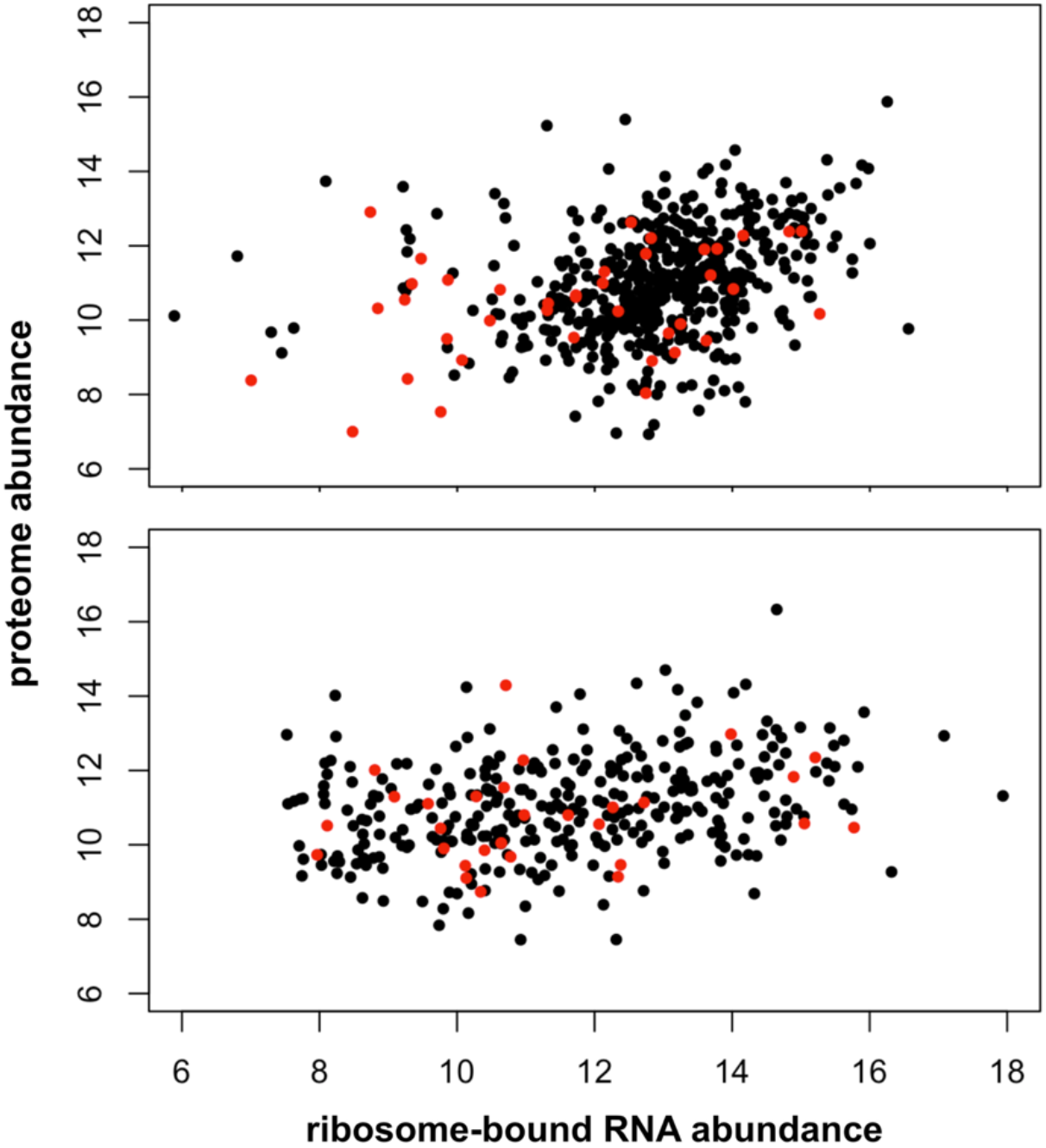
Comparison of ribosome-bound RNA levels with protein abundance in *Symbiopectobacterium endo*. (top) and *Sodalis endo*. (bottom). Abundances are shown on a normalized log2 scale, with pseudogenes colored red.

**Supplemental Figure 4:**
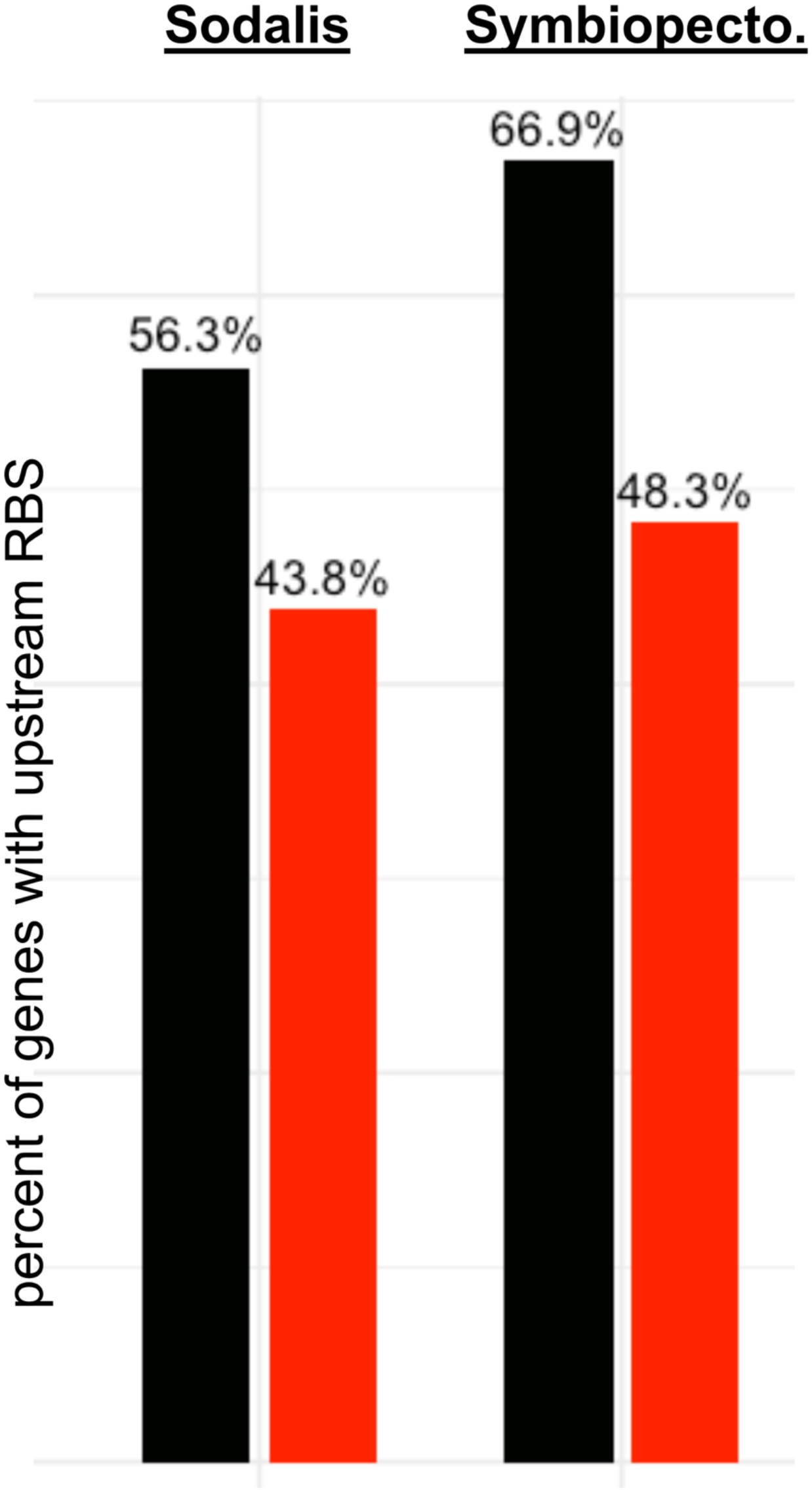
Percentage of intact (black) and pseudogenes (red) with upstream ribosomal binding sites, as predicted via Prodigal.

**Supplemental Figure 5:**
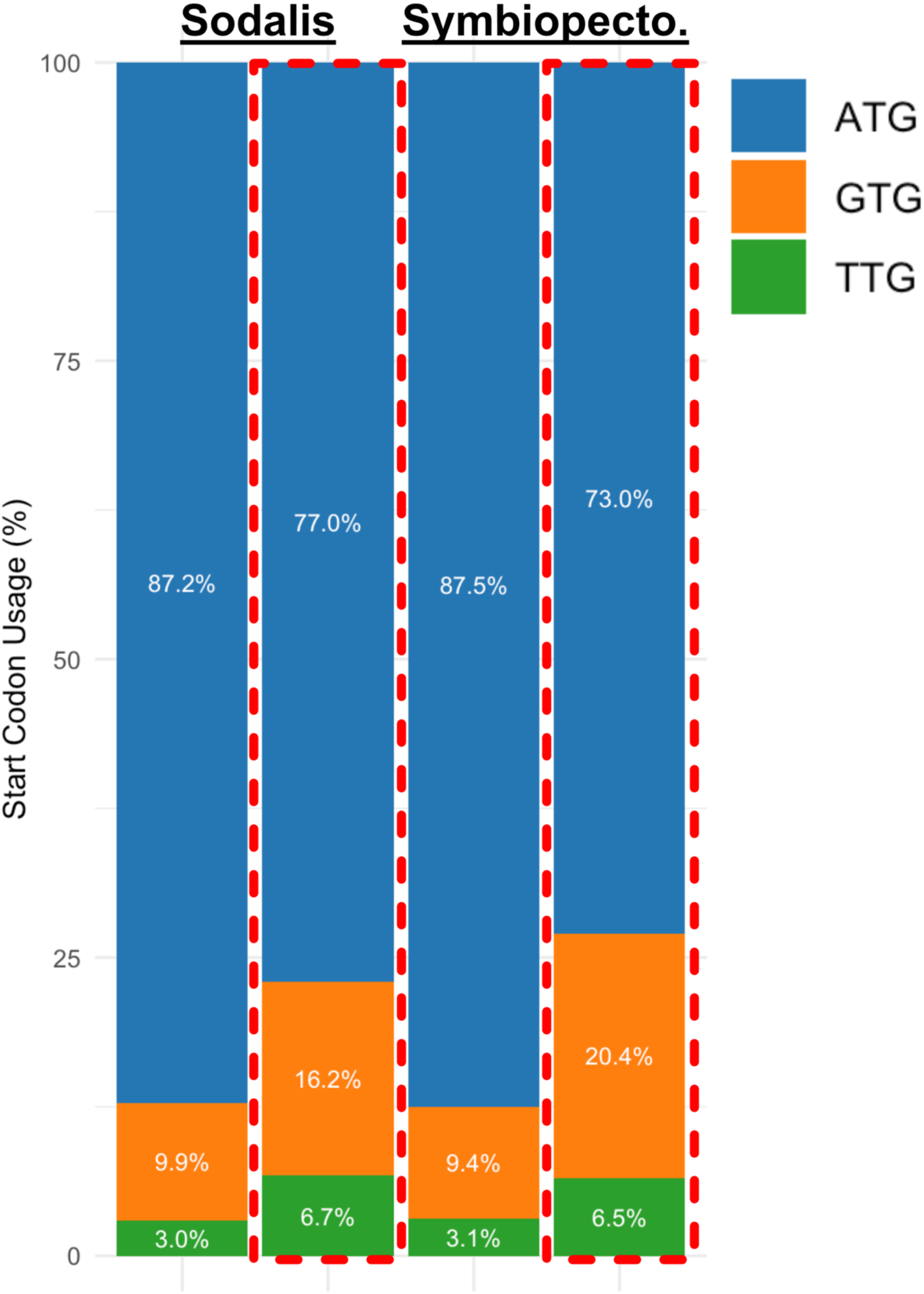
Percent of intact genes and pseudogenes (enclosed in dotted red boxes) that start with each of three types of start codons.

**Supplemental Figure 6:**
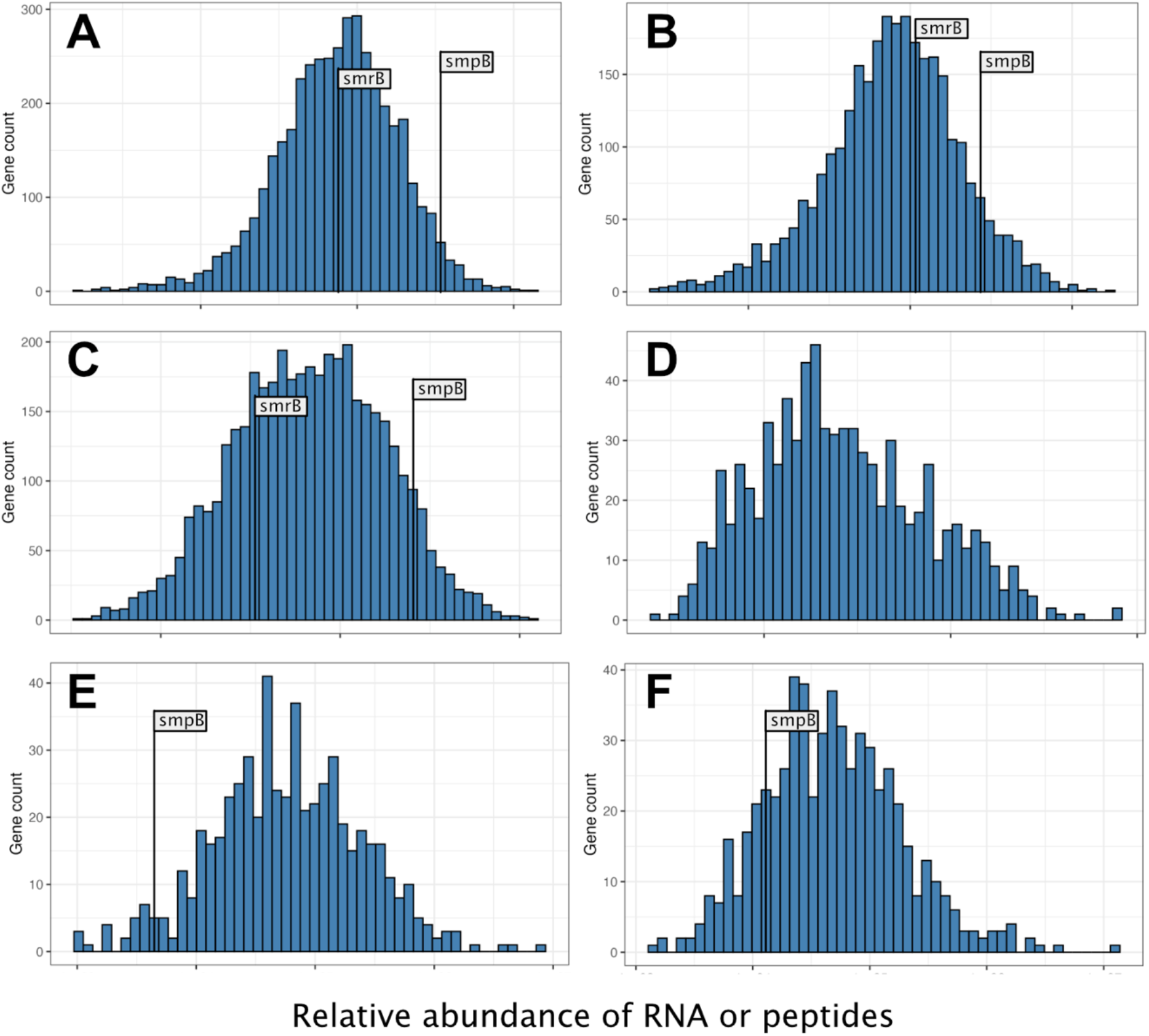
*Symbiopectobacterium endo*. and *Sodalis endo*, with an emphasis on where in each distributions the protein or transcript levels for *smpB* and *smrB* fall. Top row shows relative levels among whole transcriptomes in A) *Symbiopectobacterium* and B) *Sodalis endo*. Middle row shows relative levels among ribosome-copurified RNA in C) *Symbiopectobacterium* and D) *Sodalis endo*. Bottom row shows relative protein abundance levels in E) *Symbiopectobacterium* and F) *Sodalis endo*. Neither *smpB* nor *smrB* transcripts were detected among the *Sodalis endo*. ribosome-copurified RNA (panel D).

**Supplemental Figure 7:**
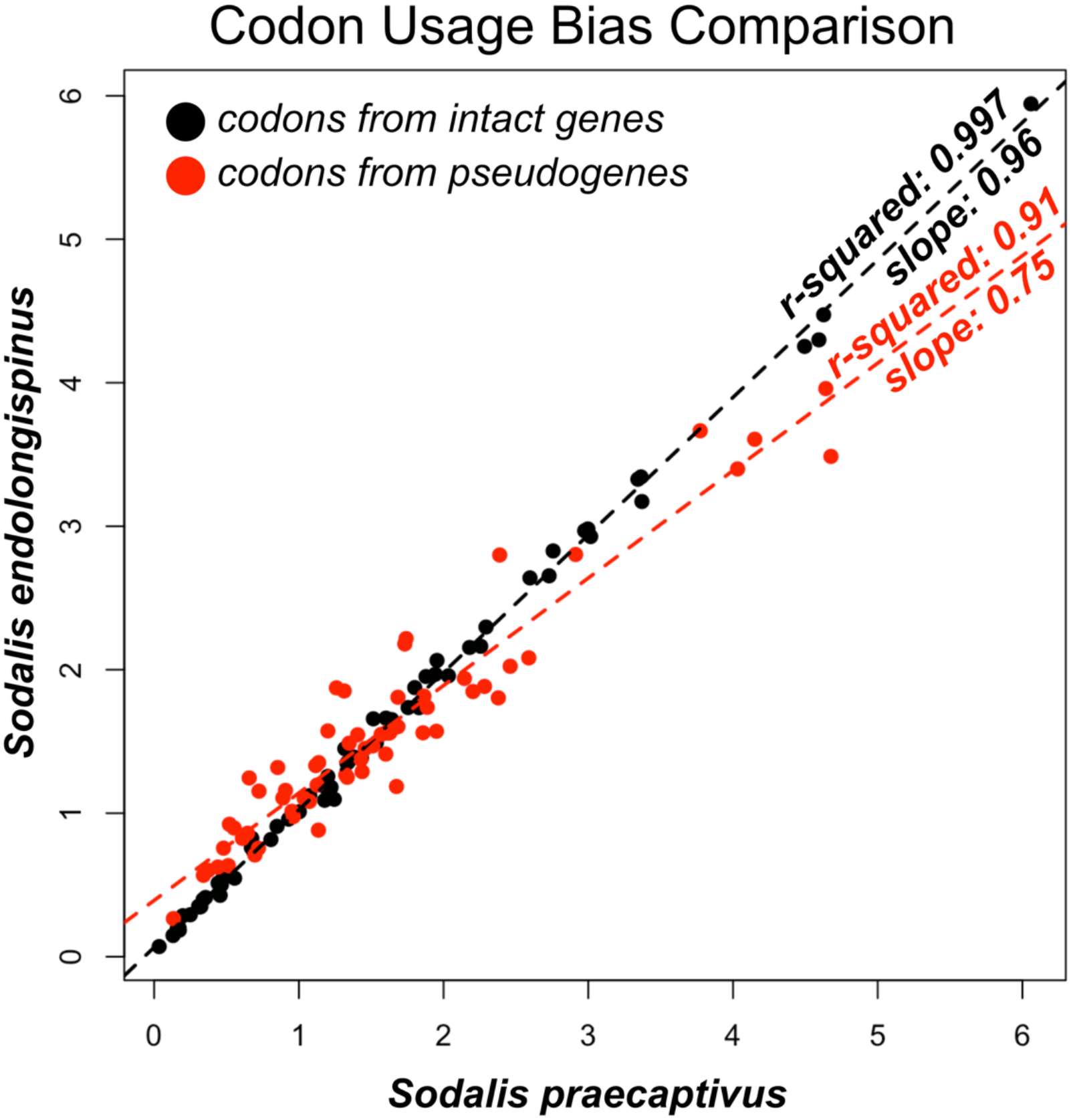
Comparison of codon usages between *Sodalis endo*. and its closest free-living relative *Sodalis praecaptivus*. Each black dot corresponds to a codon, with its frequency among the intact genes in *S. praecaptivus* shown along the x-axis and its frequency among Sodalis endo. shown along the y-axis. Codons from pseudogenes in *Sodalis endo*. are shown in red on the same scale. Linear model results superimposed over each regression.

**Supplemental Figure 8:**
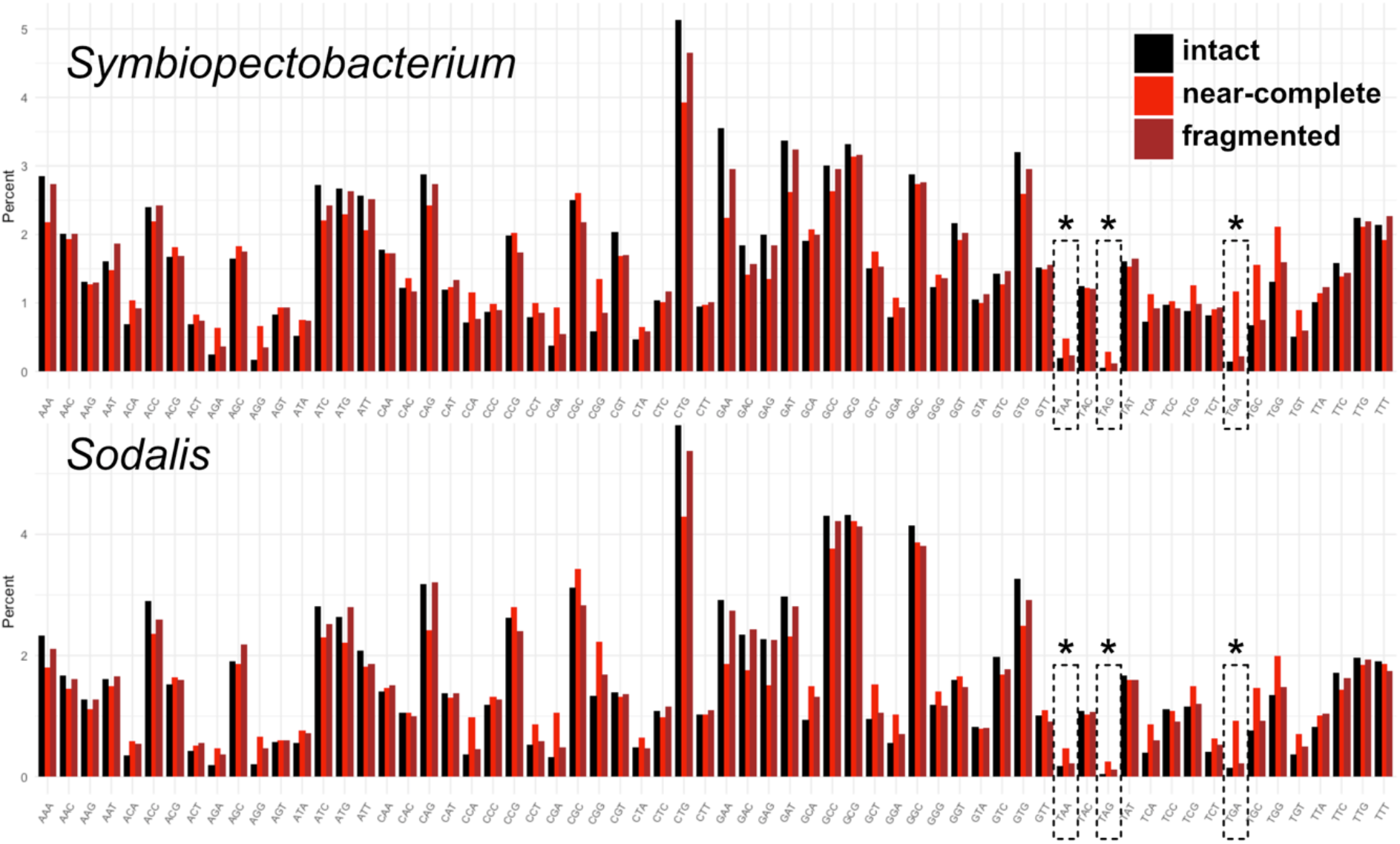
Codon usage percentages across all intact genes (black), in comparison with two types of pseudogenes: near-complete and truncated. Y-axis shows percent across all codons (within-category). Highlighted with asterisks are stop codons TAG, TGA, and TAA.

**Supplemental Figure 9:**
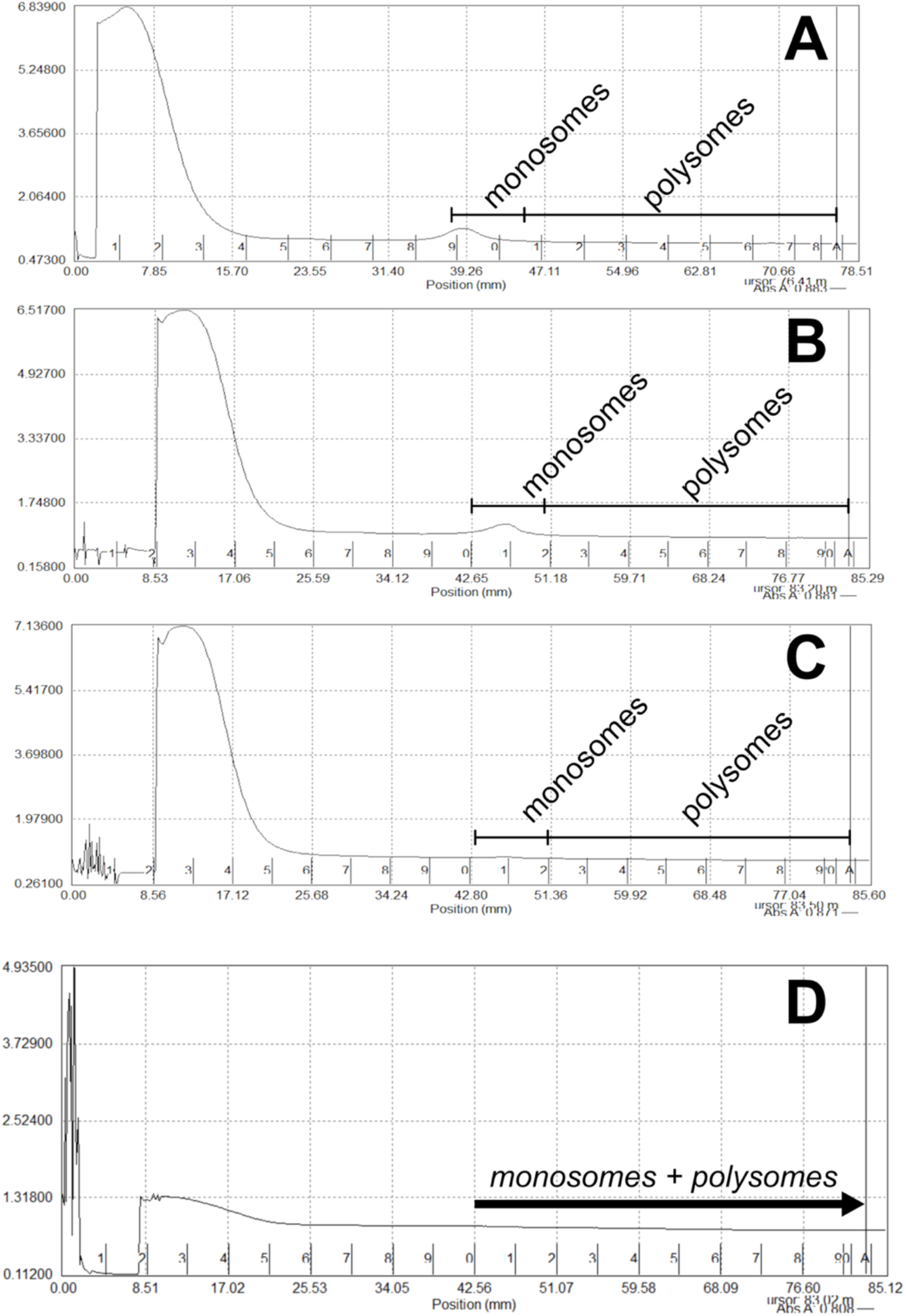
Absorbance measurements (A260), shown on the y-axis, taken during fractionation of whole-insect *Pseudococcus longispinus* samples (A-C) and P. longispinus bacteriomes (D). Fraction numbers are shown on the x-axis, with the top of the centrifuge tube corresponding to fraction 1 and the bottom (including pellet) corresponding to fraction 20. Monosome peaks are visible in panels A and B starting at fraction 10, which is the highest fraction we used for downstream sequencing and analysis (i.e., we used fractions 10-20 to infer ribosome-bound RNAs).

**Supplemental Figure 10:**
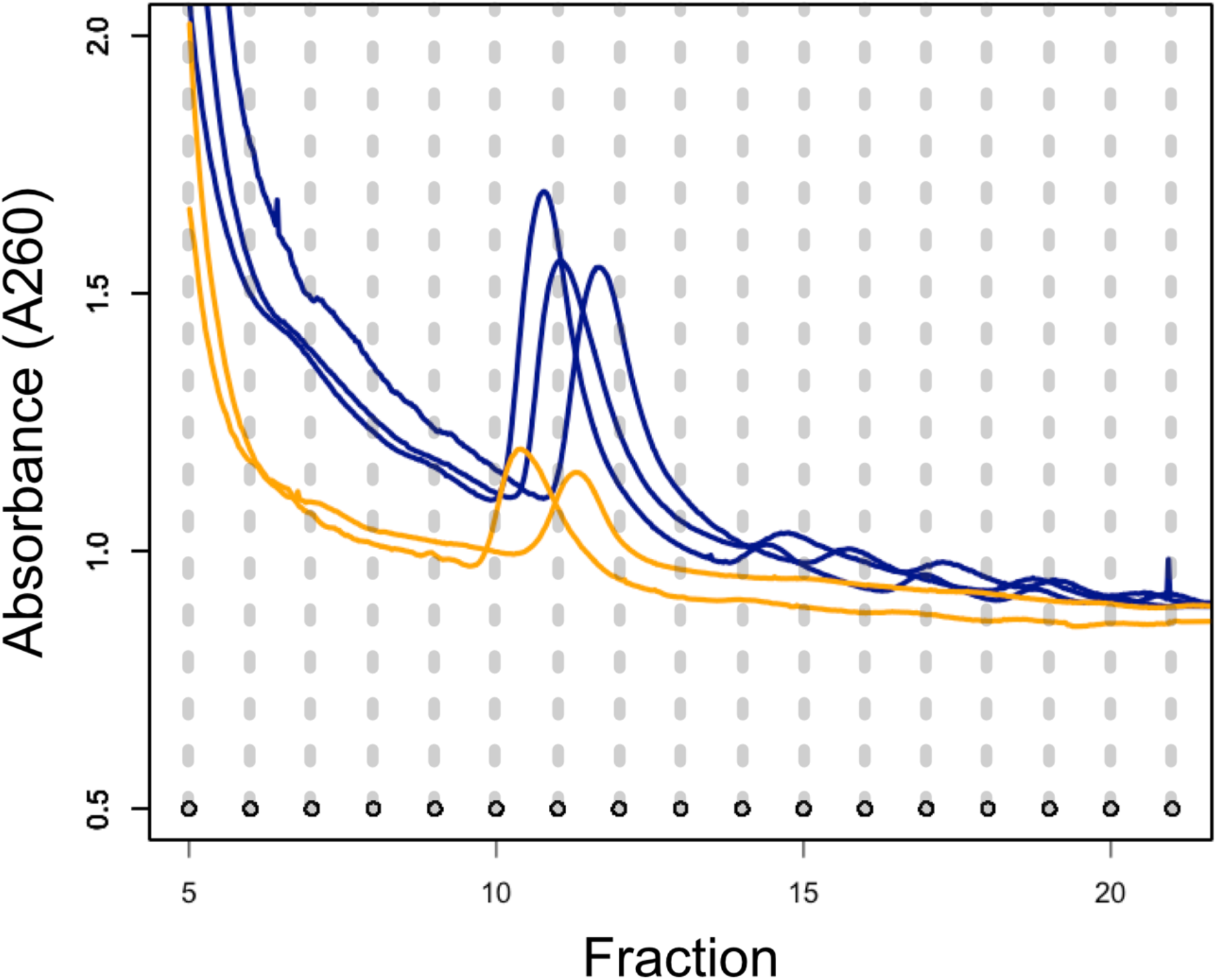
Absorbance measurements taken during fractionation of a 10-50% sucrose gradient after ultracentrifugation. Only fractions 5-20 are shown. *Sodalis praecaptivus* HS1 samples are shown in blue (triplicate), and two whole-insect *P. longispinus* samples (from panels A and B in Supplemental Figure 9) are shown in orange. Each dot on the bottom corresponds to an individual fraction from the sucrose gradient, collected automatically via the gradient fractionator.

